# Caution Regarding the Specificities of Pan-Cancer Microbial Structure

**DOI:** 10.1101/2023.01.16.523562

**Authors:** Abraham Gihawi, Colin S. Cooper, Daniel S. Brewer

**Affiliations:** Bob Champion Research & Education Building, Norwich Medical School, University of East Anglia, Norwich, UK NR4 7UQ; Earlham Institute, Norwich Research Park, Colney Lane, Norwich, UK, NR4 7UG

## Abstract

The results published in Poore and Kopylova *et al*. 2020[1] revealed the possibility of being able to almost perfectly differentiate between types of tumour based on their microbial composition using machine learning models. Whilst we believe that there is the potential for microbial composition to be used in this manner, we have concerns with the manuscript that make us question the certainty of the conclusions drawn. We believe there are issues in the areas of the contribution of contamination, handling of batch effects, false positive classifications and limitations in the machine learning approaches used. This makes it difficult to identify whether the authors have identified true biological signal and how robust these models would be in use as clinical biomarkers. We commend Poore and Kopylova *et al*. on their approach to open data and reproducibility that has enabled this analysis. We hope that this discourse assists the future development of machine learning models and hypothesis generation in microbiome research.

## Main

### Most models do not perform any better than models constructed using no information

Poore and Kopylova *et al*. detail the building of cancer type models based on microbial interrogation of TCGA (The Cancer Genome Atlas Program) cancer sequence data (which is predominantly RNA sequencing but with some whole genome sequences). Here, we evaluate these models within the framework of Whalen *et al*. 2021[2] describing common modelling pitfalls, namely: I) distributional differences, II) confounding, III) leaky preprocessing and IV) unbalanced classes.

Following their most stringent decontamination, only 5 of the 33 one-vs-all cancer type models examined were a statistically significantly improvement to models constructed using no information (at the 0.05 significance level, without false discovery correction for multiple models, “*P*-Value [Acc > NIR]”, available: http://cancermicrobiome.ucsd.edu/CancerMicrobiome_DataBrowser/) – this was not clear in the main text.

### Models pronounce nonsensical genera are informative of tumour type

Even when the model does appear to identify samples better than the negative predictor, we have concerns that many of the key features used in the model are implausible. For example, the model predicting adrenocortical carcinoma is significantly better than a negative predictor (*P*=0.002) and boasts high sensitivity (0.9565), specificity (0.998) and positive predictive values (0.71). Therefore, this model should hold some features that truly distinguish it from the remainder of cancer types. The top ten most important features for this model are Hepandensovirus (relative feature importance score: 9431, a virus that infects crustaceans[3]), *Paeniclostridium* (973), Comovirus (846), *Thalassomonas* (267, a bateria causing coral disease[4]), *Simkania* (160), *Cronobacter* (151), *Simonsiella* (148), *Leucothrix* (145, a bacteria from marine macroalgae[5]), Phikmvlikevirus (128) and N4likevirus (88). It is unclear how Phikmvlikevirus and N4likevirus might be informative for adrenocortical carcinoma as they are bacteriophages and therefore would be dependent on the co-occurrence of their bacterial hosts in the adrenal glands (or alternatively the remainder of anatomical locations[6, 7]. Many of the top performing features of other models under the most stringent decontamination approach also seem nonsensical (Table 1). This point is not covered by the Whalen pitfalls because it is generally presumed that the features being modelled exist to begin with, which in the case of taxonomic classification is not always true.

**Table 1.** Top performing features for a selection of one-vs-all cancer type models in the most stringent decontamination approach as presented Poore and Kopylova *et al*. These taxa include extremophiles that have not previously been isolated from humans. See supplementary for a full description as on NCBI of the sources for each representative species within these genera.

| Genus | Top Feature in cancer type model | Details |
| --- | --- | --- |
| *Leucothrix* | Bladder Cancer | Bacteria from marine macroalgae[5] |
| *Thalassomonas* | Uveal Melanoma | Bacteria causes disease in coral[4] |
| **Velarivirus** | Cervical Cancer | Grapevine is natural host[8] |
| **Tritimovirus** | Colon Cancer | Known to infect cereals[9] |
| **Dinovernavirus** | Renal Clear Cell Carcinoma | Contains insect viruses[10] |
| **Bacillarnavirus** | Lung Squamous Cell Carcinoma | Infects algae[11] |
| **Rymovirus** | Ovarian serous | Infects species of grass[12] |
| ***Ignicoccus*** | Prostate | Identified in marine hydrothermal vents[13] |
| ***Salinimicrobium*** | Testicular Cancer | Halophilic genus identified from marine environments[14] |

Some models do demonstrate plausible and promising results. For example, in hepatocellular carcinoma, Orthohepadnavirus is known to have a causal relationship with cancer formation[15] and has been found to be specific to the liver in other datasets [16]. This is reflected well in Poore and Kopylova *et al*.’s model where the estimated variable importance score of Orthohepadnavirus in their model (2020.53) dwarfs the next most ‘important’ feature (Levivirus, 975.09). Despite this, the model is still not significantly better than a negative predictor (*P*-value [Acc > NIR] = 1) and suffers a poor positive predictive value (0.4).

### Potential for read misclassification

We believe that these nonsensical genera arise because the models in this manuscript are built on many features that are likely to be taxonomically misclassified, from human reads or other contamination[17, 18], and therefore do not originate from microbes in the sample. One possible reason for these misclassifications is that extra steps were not taken to remove human reads prior to model building. Poore and Kopylova *et al*. detail the extraction of reads unaligned to a human reference genome which are then the subject of taxonomic classification. This pool of reads will still contain human reads which have not aligned[19]. For example, this could be because the reads are of low quality, they are mutated in cancer genomes or due to sequencing artifacts. In addition, the authors detail no human reference sequences in their taxonomic database, using 59,974 microbial genomes only. Therefore, it is highly likely that human sequences will have been misclassified as microbial. The subsequent application of SHOGUN alignment of kraken classified reads is more specific but may still involve the inappropriate classification of human reads to a database with no representation of the human genome. Additional human depletion filtering and steps to remove contamination such as those employed by the cancer microbiome atlas to distinguish tissue-resident microbiota from contaminants would have helped to remove misclassifications[20].

### Normalization introduces variance and permits modelling

Another possible contributing factor to the issues with the models is in how the data was processed. Microbiome data is dynamic[21] (Whalen I: distributional differences), and is typically heteroskedastic (meaning that the variance of a variable is non-constant over value of an independent variable *i*.*e*. the number of sequencing reads assigned to each of two genera)[22]. The authors resolve heteroskedasticity by applying a tool called Voom that is designed for RNA sequencing data of a single organism where the majority of genes have some level of expression. However, as applied by Poore *et al*. it suggests presence even when taxa are absent (Whalen III – leaky preprocessing). For example, for Hepandensovirus (genus of crustacean virus), the top feature for adrenocortical carcinoma, Voom transitions all zeros to non-zero values and untrue variation has been introduced by the global adjustment for technical variables including sequencing center (figure 1**a**, batch correction relating to Whalen II: confounding). Therefore, this normalization appears beneficial on the global level but raises prominent concerns at the level of individual taxa.

**Figure 1:**
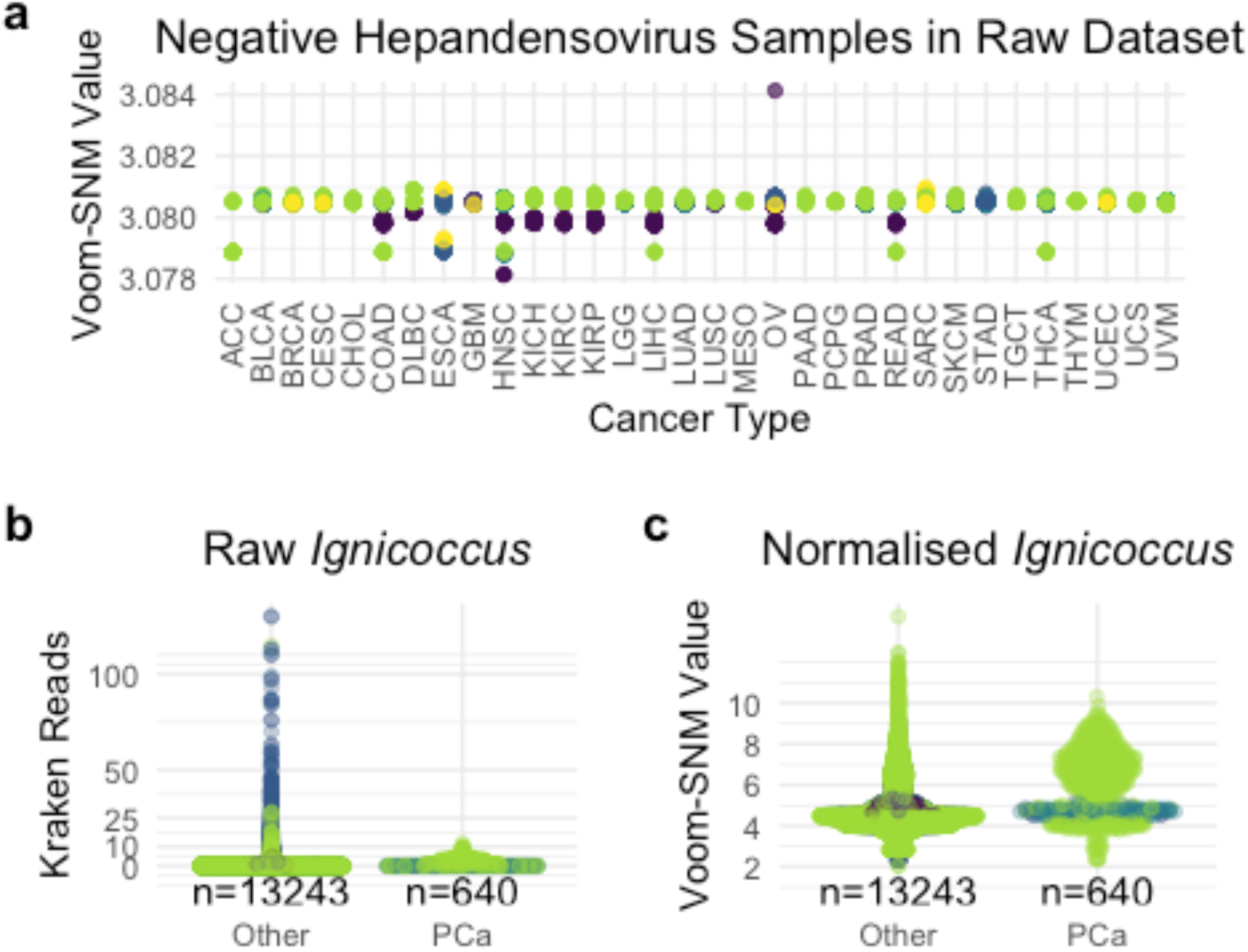
**a)** Voom-SNM normalised TCGA samples (n = 17,624) that were negative for crustacean virus hepandensovirus with zero classified reads in the original Kraken dataset with the most stringent decontamination approach. One sample contained two sequencing reads for Hepandensovirus which has been omitted from this figure to illustrate inappropriate variation introduced by SNM. The colour of each point indicates the sequencing center. The x-axis demonstrates cancer types using TCGA abbreviations as in Poore et al.[1]. Raw **(b)** and Voom-SNM normalised **(c)** Ignicoccus values which was deemed the most important feature for predicting prostate cancer (PCa) from all other cancer types (n=13,883 primary tumours). Median values are as follows: Kraken raw other 0, Kraken raw PCa 1, normalised other 4.49, normalised PCa 5.82. In both the raw and normalised cases, the distributions are significantly different (Wilcox signed rank-sum test P < 2.2 × 10^−16^)

Another example of how the processing of data can be problematic is provided by the extremophile genera *Ignicoccus* in prostate cancer samples. *Ignicoccus* demonstrates a statistically significant increase in prostate cancer samples compared to other cancers in the normalized dataset (Wilcox signed rank-sum test *P*<2.2 × 10^−16^, figure 1**b-c**). In the raw, unprocessed data no increase in prostate cancer samples is apparent. Indeed, most values are zero and the maximum number of reads found in the raw prostate cancer data for *Ignicoccus* is 12 (low evidence of detection). It is also highly likely that these are false taxonomic assignments given that *Ignicoccus* was identified in marine hydrothermal vents[13]. This taxon should have been filtered out prior to model building – the application of a minimum read threshold (*i*.*e*. 100 classified reads) would have assisted the removal of spurious taxa.

### The models are trained on unbalanced data

The performance of the models may in part be due to the major imbalance in class size in the datasets (Whalen IV: unbalanced classes); meaning that before model construction, data in the cancer set under investigation are multiplied up many times (upsampling) so that patient numbers in the “cancer groups” and in the “all other cancers group” become similar. This approach may amplify the prominence of implausible artifactual data. Adrenocortical carcinoma for example has 79 associated samples (as per Metadata-TCGA-All-18116-Samples.csv provided by Poore *et al*.). This means that 18,037 are not adrenocortical carcinoma. Adrenocortical carcinoma therefore represents 0.44% of the whole dataset and therefore data from adrenocortical carcinoma is amplified up to 230 times to equal the sample size of the rest of the dataset. The modelling is therefore overexposed to inappropriate variation in taxa such as Hepandensovirus (figure 1).

### Discussion

Ideally, the authors would have followed the RIDE criteria set out by Eisenhofer *et al*. (also authored by Knight) as closely as possible[23]. Some of these criteria are difficult, if not impossible to meet on this dataset, but this means that the context of this study should be in hypothesis generation and more care should be taken in the conclusions drawn. Poore and Kopylova *et al*. have paid considerable attention to the issue of contamination but nonsensical taxa with limited evidence of true involvement are still prominent. The hypothesis that the tumour microbiome is dependent on the anatomical site is well founded based on prior work[24], but the models produced by Poore and Kopylova *et al*. are at best suggestive and do not substantiate this observation.

Poore and Kopylova *et al*. use many good practices in machine learning[25] but there is the need to avoid the pitfalls of Whalen *et al*. and use more stringent methods regarding contamination, taxonomic misclassification and a lack of explanations backed by microbiological insights for approximately 600 models. It is advisable that additional care should be taken to include only taxa with strong evidence of presence based on computational evidence, consideration of the likelihood of contamination and prior biological evidence that the taxa exist in the biological sample of interest.

## Conclusion

We believe that the tumour microbiome is an exciting field and that using large sequencing datasets with rich metadata may unlock much more about the nature of the interplay between microbes and cancer. The authors have stated that the tumour microbiome is specific to the tumour type using machine learning models, but we have concerns. There needs to be a more appropriate demonstration of microbial differences between tumour types and stringent validation of models before we can be certain of these differences. This is required before these findings can be translated into the clinic. A dataset with a less prominent batch effect, more balanced class sizes, modelling tumour type (not one-vs-all models) might help to better distinguish pan-cancer microbial structure.

## Methods

All analysis in this manuscript was conducted on the open-source data made available by Poore and Kopylova *et al*.*[1]* available at: ftp.microbio.me/pub/cancer_microbiome_analysis/. Files analysed include: Kraken-TCGA-Raw-Data-17625-Samples.csv (MD5 checksum: 6af81818f69bf56b79836e1c317c3e03), Metadata-TCGA-All-18116-Samples.csv (MD5 checksum: dbdd1f64d45973977fc8435db2eb8b3e), Kraken-TCGA-Voom-SNM-Most-Stringent-Filtering-Data.csv (MD5 checksum: b7e50700b791b8881426aeb1fa12c3bb).

Model performance and feature importance was accessed: http://cancermicrobiome.ucsd.edu/CancerMicrobiome_DataBrowser/. All data was analysed in R (version 4.2.1). Packages used include tidyverse[26] (version 1.3.2), ggpubr[27] (version 0.5.0), ggbeeswarm[28] (version 0.7.1), cowplot[29] (version 1.1.1) and EnvStats[30] (version 2.7.0). Hypothesis testing was performed with the wilcox.test() function.

Representative species within top features (supplementary) were identified by browsing GTDB[31] (release version 207). Associated metadata regarding isolation sources was found by accessing links presented on the GTDB taxonomy browser.

## References

[1] Poore GD, Kopylova E, Zhu Q, Carpenter C, Fraraccio S, Wandro S, et al. Microbiome analyses of blood and tissues suggest cancer diagnostic approach. Nature. 2020;579:567–74.

[2] Whalen S, Schreiber J, Noble WS, Pollard KS. Navigating the pitfalls of applying machine learning in genomics. Nat Rev Genet. 2021.

[3] Cotmore SF, Agbandje-McKenna M, Chiorini JA, Mukha DV, Pintel DJ, Qiu J, et al. The family Parvoviridae. Arch Virol. 2014;159:1239–47.

[4] Hosoya S, Adachi K, Kasai H. Thalassomonas actiniarum sp. nov. and Thalassomonas haliotis sp. nov., isolated from marine animals. Int J Syst Evol Microbiol. 2009;59:686–90.

[5] Liu T, Zhang Y, Zhang X, Zhou L, Meng C, Zhou C, et al. Leucothrix sargassi sp. nov., isolated from a marine alga [Sargassum natans (L.) Gaillon]. Int J Syst Evol Microbiol. 2019;69:3857–62.

[6] Wittmann J, Klumpp J, Moreno Switt AI, Yagubi A, Ackermann HW, Wiedmann M, et al. Taxonomic reassessment of N4-like viruses using comparative genomics and proteomics suggests a new subfamily - “Enquartavirinae”. Arch Virol. 2015;160:3053–62.

[7] Merabishvili M, Vandenheuvel D, Kropinski AM, Mast J, De Vos D, Verbeken G, et al. Characterization of newly isolated lytic bacteriophages active against Acinetobacter baumannii. PLoS One. 2014;9:e104853.

[8] Yu H, Qi S, Chang Z, Rong Q, Akinyemi IA, Wu Q. Complete genome sequence of a novel velarivirus infecting areca palm in China. Arch Virol. 2015;160:2367–70.

[9] Rabenstein F, Seifers DL, Schubert J, French R, Stenger DC. Phylogenetic relationships, strain diversity and biogeography of tritimoviruses. J Gen Virol. 2002;83:895–906.

[10] Roundy CM, Azar SR, Rossi SL, Weaver SC, Vasilakis N. Insect-Specific Viruses: A Historical Overview and Recent Developments. Adv Virus Res. 2017;98:119–46.

[11] Short SM, Staniewski MA, Chaban YV, Long AM, Wang D. Diversity of Viruses Infecting Eukaryotic Algae. Curr Issues Mol Biol. 2020;39:29–62.

[12] Webster DE, Beck DL, Rabenstein F, Forster RL, Guy PL. An improved polyclonal antiserum for detecting Ryegrass mosaic rymovirus. Arch Virol. 2005;150:1921–6.

[13] Paper W, Jahn U, Hohn MJ, Kronner M, Nather DJ, Burghardt T, et al. Ignicoccus hospitalis sp. nov., the host of ‘Nanoarchaeum equitans’. Int J Syst Evol Microbiol. 2007;57:803–8.

[14] Nedashkovskaya OI, Vancanneyt M, Kim SB, Han J, Zhukova NV, Shevchenko LS. Salinimicrobium marinum sp. nov., a halophilic bacterium of the family Flavobacteriaceae, and emended descriptions of the genus Salinimicrobium and Salinimicrobium catena. Int J Syst Evol Microbiol. 2010;60:2303–6.

[15] Ringelhan M, McKeating JA, Protzer U. Viral hepatitis and liver cancer. Philos Trans R Soc Lond B Biol Sci. 2017;372.

[16] Zapatka M, Borozan I, Brewer DS, Iskar M, Grundhoff A, Alawi M, et al. The landscape of viral associations in human cancers. Nat Genet. 2020;52:320–30.

[17] Salter SJ, Cox MJ, Turek EM, Calus ST, Cookson WO, Moffatt MF, et al. Reagent and laboratory contamination can critically impact sequence-based microbiome analyses. BMC Biology. 2014;12:87.

[18] de Goffau MC, Lager S, Sovio U, Gaccioli F, Cook E, Peacock SJ, et al. Human placenta has no microbiome but can contain potential pathogens. Nature. 2019;572:329–34.

[19] Gihawi A, Rallapalli G, Hurst R, Cooper CS, Leggett RM, Brewer DS. SEPATH: benchmarking the search for pathogens in human tissue whole genome sequence data leads to template pipelines. Genome Biol. 2019;20:208.

[20] Dohlman AB, Arguijo Mendoza D, Ding S, Gao M, Dressman H, Iliev ID, et al. The cancer microbiome atlas: a pan-cancer comparative analysis to distinguish tissue-resident microbiota from contaminants. Cell Host Microbe. 2021;29:281–98 e5.

[21] Gerber GK. The dynamic microbiome. FEBS Lett. 2014;588:4131–9.

[22] McMurdie PJ. Normalization of Microbiome Profiling Data. Methods Mol Biol. 2018;1849:143–68.

[23] Eisenhofer R, Minich JJ, Marotz C, Cooper A, Knight R, Weyrich LS. Contamination in Low Microbial Biomass Microbiome Studies: Issues and Recommendations. Trends Microbiol. 2019;27:105–17.

[24] Costello EK, Lauber CL, Hamady M, Fierer N, Gordon JI, Knight R. Bacterial community variation in human body habitats across space and time. Science. 2009;326:1694–7.

[25] Knight R, Vrbanac A, Taylor BC, Aksenov A, Callewaert C, Debelius J, et al. Best practices for analysing microbiomes. Nat Rev Microbiol. 2018;16:410–22.

[26] Wickham H, Averick M, Bryan J, Chang W, McGowan L, François R, et al. Welcome to the tidyverse. Journal of Open Source Software. 2019;4:1686.

[27] Kassambara A. ggpubr: ‘ggplot2’ Based Publication Ready Plots. 2022.

[28] Clarke ES-MSD, C. ggbeeswarm: Categorical Scatter (Violin Point) Plots. 2022.

[29] Wilke C. cowplot: Streamlined Plot Theme and Plot Annotations for ‘ggplot2’. 2020.

[30] Millard S. EnvStats: An R Package for Environmental Statistics: Springer; 2013.

[31] Parks DH, Chuvochina M, Rinke C, Mussig AJ, Chaumeil PA, Hugenholtz P. GTDB: an ongoing census of bacterial and archaeal diversity through a phylogenetically consistent, rank normalized and complete genome-based taxonomy. Nucleic Acids Res. 2022;50:D785–D94.

