## Supplementary Data for "Caution Regarding the Specificities of Pan-Cancer Microbial Structure"

Supplementary Table 1 - These species were found on GTDB taxonomy and include species representatives that passed quality checks. Each of these species originates from a genus that was the top performing microbial feature in the one-vs-all tumour models provided by Poore *et al*. [1] and was considered the most stringent approach to decontamination.

| **GTDB Taxonomy** | **Isolation Source Information** |
| --- | --- |
| Leucothrix sp002747455 | swab sample of gingival sulcus (mouth) from 29 year old lactating female Dolphin_Z <https://www.ncbi.nlm.nih.gov/biosample/SAMN07757601/> |
| Leucothrix mucor | seaweed, Friday Harbor, WA <https://www.ncbi.nlm.nih.gov/biosample/SAMN02440559/> |
| Leucothrix pacifica | Seawater - [Pacific Ocean: South Pacific Gyre](https://www.ncbi.nlm.nih.gov/biosample?term=%22geo_loc_name=Pacific%20Ocean:%20South%20Pacific%20Gyre%22%5battr%5d)  <https://www.ncbi.nlm.nih.gov/biosample/SAMN09228920/> |
| Leucothrix arctica | Seawater - [Svalbard: coast of Kongsfjorden, Arctic](https://www.ncbi.nlm.nih.gov/biosample?term=%22geo_loc_name=Svalbard:%20coast%20of%20Kongsfjorden,%20Arctic%22%5battr%5d)  <https://www.ncbi.nlm.nih.gov/biosample/SAMN09228919/> |
| Thalassomonas actiniarum | Sea anemone  <https://www.ncbi.nlm.nih.gov/biosample/SAMN03278135/> |
| Thalassomonas viridans | Cultivated oysters  <https://www.ncbi.nlm.nih.gov/biosample/SAMN03278134/> |
| Ignicoccus_A sp013154085 | deepsea hydrothermal sulfide chimney  <https://www.ncbi.nlm.nih.gov/biosample/SAMN12405658/> |
| Ignicoccus_A sp015521275 | marine hydrothermal vent biome  <https://www.ncbi.nlm.nih.gov/biosample/SAMN12837294/> |
| Ignicoccus_A hospitalis | ***Ignicoccus* hospitalis Kin4/I**. *Ignicoccus* hospitalis Kin4/I was isolated from gravel obtained from the shallow marine hydrothermal system of the Kolbeinsey Ridge, north of Iceland and will be used for comparative analysis with other thermophiles  <https://www.ncbi.nlm.nih.gov/bioproject/13914> |
| Ignicoccus islandicus | “The genome sequence of this strain will provide information on the adaptation to high temperature environments and be used for evolutionary and comparative genomic analysis.”  <https://www.ncbi.nlm.nih.gov/bioproject/PRJNA61347/> |
| Salinimicrobium xinjiangense | Soil sediment of a salt lake  <https://www.ncbi.nlm.nih.gov/biosample/SAMN02440840/> |
| Salinimicrobium terrae | Saline soil  <https://www.ncbi.nlm.nih.gov/biosample/SAMN02440676/> |
| Salinimicrobium sp012038135 | marine sediment  https://www.ncbi.nlm.nih.gov/biosample/SAMN14480397/ |
| Salinimicrobium marinum | Unstated <https://www.ncbi.nlm.nih.gov/biosample/SAMD00245570/> |
| Salinimicrobium catena | Unstated <https://www.ncbi.nlm.nih.gov/biosample/SAMN04488140/> |
| Salinimicrobium sediminis | Unstated <https://www.ncbi.nlm.nih.gov/biosample/SAMN06296241/> |

Supplementary Table 2 – Viruses as top features in one-vs-all models. These were identified by searching NCBI at genus level and investigating each individual species. Each of these species originates from a genus that was the top performing feature in one of the one-vs-all tumour models presented by Poore *et al*. under the most stringent decontamination conditions.

| [*Areca palm velarivirus 1*](https://www.ncbi.nlm.nih.gov/data-hub/taxonomy/1654603) | Complete genome sequence of a novel Velarivirus infecting areca palm in China  <https://www.ncbi.nlm.nih.gov/nuccore/NC_027121.1/> |
| --- | --- |
| [*Cordyline virus 1*](https://www.ncbi.nlm.nih.gov/data-hub/taxonomy/937809) (velarivirus) | An assemblage of closteroviruses infects Hawaiian ti (Cordyline fruticosa L.) <https://www.ncbi.nlm.nih.gov/nuccore/NC_038421.1/> |
| *Cordyline virus* 2 (velarivirus) | Differentiation, distribution, and elimination of closteroviruses infecting Cordyline fruticosa (L.) in Hawaii  <https://www.ncbi.nlm.nih.gov/nuccore/NC_043453.1/> |
| [*Cordyline virus 3*](https://www.ncbi.nlm.nih.gov/data-hub/taxonomy/1177752) (velarivirus) | Differentiation, distribution, and elimination of closteroviruses infecting Cordyline fruticosa (L.) in Hawaii  <https://www.ncbi.nlm.nih.gov/nuccore/NC_043107.1/> |
| [*Cordyline virus 4*](https://www.ncbi.nlm.nih.gov/data-hub/taxonomy/1177753) (velarivirus) | Differentiation, distribution, and elimination of closteroviruses infecting Cordyline fruticosa (L.) in Hawaii <https://www.ncbi.nlm.nih.gov/nuccore/NC_043108.1/> |
| [*Grapevine leafroll-associated virus 7*](https://www.ncbi.nlm.nih.gov/data-hub/taxonomy/217615) (velarivirus) | Molecular characterization and taxonomy of grapevine leafroll-associated virus 7 <https://www.ncbi.nlm.nih.gov/nuccore/NC_016436.1/> |
| [*Little cherry virus 1*](https://www.ncbi.nlm.nih.gov/data-hub/taxonomy/217686)  [*Malus domestica virus A*](https://www.ncbi.nlm.nih.gov/data-hub/taxonomy/2664236) (velarivirus) | Complete genome structure and phylogenetic analysis of little cherry virus, a mealybug-transmissible closterovirus  <https://www.ncbi.nlm.nih.gov/nuccore/NC_001836.1/> |
| [*Malus domestica virus A*](https://www.ncbi.nlm.nih.gov/data-hub/taxonomy/2664236)  (valarivirus) | Genomic characterization of a novel velarivirus Malus domestica virus A (MdoVA) infecting apple  <https://www.ncbi.nlm.nih.gov/nuccore/NC_055599.1/> |
| [*Brome streak mosaic virus*](https://www.ncbi.nlm.nih.gov/data-hub/taxonomy/42631)  (***[Tritimovirus](https://www.ncbi.nlm.nih.gov/data-hub/taxonomy/156207)***) | The complete nucleotide sequence and genome organization of the mite-transmitted brome streak mosaic rymovirus in comparison with those of potyviruses <https://www.ncbi.nlm.nih.gov/nuccore/NC_003501.1/> |
| [*Oat necrotic mottle virus*](https://www.ncbi.nlm.nih.gov/data-hub/taxonomy/112437)  (***[Tritimovirus](https://www.ncbi.nlm.nih.gov/data-hub/taxonomy/156207)***) | Complete nucleotide sequence of Oat necrotic mottle virus: a distinct Tritimovirus species (family Potyviridae) most closely related to Wheat streak mosaic virus  <https://www.ncbi.nlm.nih.gov/nuccore/NC_005136.1/> |
| [*Tall oatgrass mosaic virus*](https://www.ncbi.nlm.nih.gov/data-hub/taxonomy/1414644)  (***[Tritimovirus](https://www.ncbi.nlm.nih.gov/data-hub/taxonomy/156207)***) | Tall oatgrass mosaic virus (TOgMV): a novel member of the genus Tritimovirus infecting Arrhenatherum elatius <https://www.ncbi.nlm.nih.gov/nuccore/NC_022745.1/> |
| [*Wheat eqlid mosaic virus*](https://www.ncbi.nlm.nih.gov/data-hub/taxonomy/460364)  (***[Tritimovirus](https://www.ncbi.nlm.nih.gov/data-hub/taxonomy/156207)***) | Analyses of the complete sequence of the genome of wheat Eqlid mosaic virus, a novel species in the genus Tritimovirus  <https://www.ncbi.nlm.nih.gov/nuccore/NC_009805.1/> |
| [*Wheat streak mosaic virus*](https://www.ncbi.nlm.nih.gov/data-hub/taxonomy/31741)  (***[Tritimovirus](https://www.ncbi.nlm.nih.gov/data-hub/taxonomy/156207)***) | Phylogenetic relationships within the family potyviridae: wheat streak mosaic virus and brome streak mosaic virus are not members of the genus rymovirus <https://www.ncbi.nlm.nih.gov/nuccore/NC_001886.1/> |
| [*Yellow oat grass mosaic virus*](https://www.ncbi.nlm.nih.gov/data-hub/taxonomy/2170240)  (***[Tritimovirus](https://www.ncbi.nlm.nih.gov/data-hub/taxonomy/156207)***) | Genome sequence of two isolates of Yellow oatgrass mosaic virus, a new grass-infecting Tritimovirus <https://www.ncbi.nlm.nih.gov/nuccore/NC_024471.1/> |
| [*Aedes pseudoscutellaris reovirus*](https://www.ncbi.nlm.nih.gov/data-hub/taxonomy/341721) (Dinovernavirus) | isolated from Aedes pseudoscutellaris mosquito cells  <https://www.ncbi.nlm.nih.gov/nuccore/NC_007666.1/>  10.1016/j.virol.2005.08.028 |
| [*Chaetoceros socialis forma radians RNA virus 1*](https://www.ncbi.nlm.nih.gov/data-hub/taxonomy/2169725)  (Bacillarnavirus) | Isolation and characterization of a single-stranded RNA virus infecting the bloom-forming diatom Chaetoceros socialis <https://www.ncbi.nlm.nih.gov/nuccore/NC_012212.1/> |
| [*Chaetoceros tenuissimus RNA virus 01*](https://www.ncbi.nlm.nih.gov/data-hub/taxonomy/497136) (Bacillarnavirus) | Isolation and characterization of a single-stranded RNA virus infecting the marine planktonic diatom Chaetoceros tenuissimus Meunier <https://www.ncbi.nlm.nih.gov/nuccore/NC_038321.1/> |
| [*Rhizosolenia setigera RNA virus 01*](https://www.ncbi.nlm.nih.gov/data-hub/taxonomy/359987) (Bacillarnavirus) | Complete nucleotide sequence of a single-stranded RNA virus infecting the bloom-forming diatom Rhizosolenia setigera <https://www.ncbi.nlm.nih.gov/nuccore/NC_018613.1/> |
| [*Agropyron mosaic virus*](https://www.ncbi.nlm.nih.gov/data-hub/taxonomy/41763)  (Rymovirus) | Functional replacement of Wheat streak mosaic virus HC-Pro with the corresponding cistron from a diverse array of viruses in the family  <https://www.ncbi.nlm.nih.gov/nuccore/NC_005903.1/> |
| [*Hordeum mosaic virus*](https://www.ncbi.nlm.nih.gov/data-hub/taxonomy/41764)  (Rymovirus) | Functional replacement of Wheat streak mosaic virus HC-Pro with the corresponding cistron from a diverse array of viruses in the family Potyviridae  <https://www.ncbi.nlm.nih.gov/nuccore/NC_005904.1/> |
| [*Ryegrass mosaic virus*](https://www.ncbi.nlm.nih.gov/data-hub/taxonomy/40666)  (Rymovirus) | The complete nucleotide sequence of the Ryegrass mosaic virus  <https://www.ncbi.nlm.nih.gov/nuccore/NC_001814.1/> |

[1] Poore GD, Kopylova E, Zhu Q, Carpenter C, Fraraccio S, Wandro S, et al. Microbiome analyses of blood and tissues suggest cancer diagnostic approach. Nature. 2020;579:567-74.
